# Draft genome sequence of the *Solanum aethiopicum* provides insights into disease resistance, drought tolerance and the evolution of the genome

**DOI:** 10.1101/532077

**Authors:** Bo Song, Yue Song, Yuan Fu, Elizabeth Balyejusa Kizito, Pamela Nahamya Kabod, Huan Liu, Sandra Ndagire Kamenya, Samuel Muthemba, Robert Kariba, Xiuli Li, Sibo Wang, Shifeng Cheng, Alice Muchugi, Ramni Jamnadass, Howard-Yana Shapiro, Allen Van Deynze, Huanming Yang, Jian Wang, Xun Xu, Damaris Achieng Odeny, Xin Liu

## Abstract

**Background:** *S. aethiopicum* is a close relative to *S. melongena* and has been routinely used to improve disease resistance in *S. melongena*. However, these efforts have been greatly limited by the lack of a reference genome and the clear understanding of the genes involved during biotic and abiotic stress response.

**Results:** We present here a draft genome assembly of *S. aethiopicum* of 1.02 Gb in size, which is predominantly occupied by repetitive sequences (76.2%), particularly long terminal repeat elements. We annotated 37,681 gene models including 34,905 protein-coding genes. We observed an expansion of resistance genes through two rounds of amplification of LTR-Rs, occurred around 1.25 and 3.5 million years ago, respectively. The expansion also occurred in gene families related to drought tolerance. A number of 14,995,740 SNPs are identified by re-sequencing 65 *S. aethiopicum* genotypes including “Gilo” and “Shum” accessions, 41,046 of which are closely linked to resistance genes. The domestication and demographic history analysis reveals selection of genes involved in drought tolerance in both “Gilo” and “Shum” groups. A pan-genome of *S. aethiopicum* with a total of 36,250 protein-coding genes was assembled, of which 1,345 genes are missing in the reference genome.

**Conclusions:** Overall, the genome sequence of *S. aethiopicum* increases our understanding of the genomic mechanisms of its extraordinary disease resistance and drought tolerance. The SNPs identified are available for potential use by breeders. The information provided here will greatly accelerate the selection and breeding of the African eggplant as well as other crops within the Solanaceae family.

## Background

African eggplant, *Solanum aethiopicum*, is an indigenous non-tuberiferous *Solanum* crop, which is majorly grown in tropical Africa [1] and is important in Central and West Africa. The fruits and leaves are eaten fresh or cooked. It is reported to have medicinal value and its roots and fruits have been used to treat colic, high blood pressure and uterine complications in Africa (FAO). Experiments performed in rats showed that the methanol extract of *S. aethiopicum* has anti-inflammatory activity [2]. *S. aethiopicum* is generally classified into four groups based on the use; Gilo, Shum, Kumba and Aculeatum. As the most important group, Gilo has edible fruits, while Shum has small and bitter fruits; Kumba is usually used as a leafy vegetable; Aculeatum is used as rootstocks due to its excellent disease resistance nature (mansfeld.ipk-gatersleben.de).

Although *S. aethiopicum* is the second most cultivated eggplant, as an “orphan crop”, research and breeding investments are substantially lagging behind, compared to other Solanaceae relatives such as tomato, potato and eggplant, partly because of the lack of a reference genome sequence. Genomics-assisted breeding is an effective approach advancing the breeding of orphan crops. Attempts have been made to develop molecular markers for *S. aethiopicum* using *S. melongena* genome as a reference, but with a cost of compromising accuracy [3]. Another approach involves genome editing. Lemmon et al. (2018) mutated or pruned genes according to the structures of their orthologues in domesticated tomato, and rapidly improved three major productivity traits of a Solanaceae crop, *Physalis pruinose* using clustered regularly interspaced short palindromic repeats (CRISPR)–CRISPR-associated protein-9 nuclease (Cas9) (CRISPR–Cas9) [4]. Both approaches rely on the availability of genome and gene sequences.

Orphan crops also serve as gene reservoirs which are valuable to their crop relatives. For example, due to its cross-compatibility with *S. melongena* [5, 6] and its outstanding resistance to various pathogens including *Fusarium, Ralstonia* and *Verticillium* [7-9], *S. aethiopicum* has been used to develop rootstocks [9] or improve the disease resistance of *S. melongena* [10]. As the genomic basis of resistance in *S. aethiopicum* is poorly understood, resistance improvement through interspecies cross can be time consuming. This challenge could be overcome by locating resistance genes and developing markers associated with the respective genes. The birth and expansion of resistance genes are usually accompanied with the amplification of LTR-Rs. A typical example is shown in hot pepper (*Capsicum annuum*), also a Solanaceous vegetable. The burst of LTR-Rs has substantially mediated the retrotransposition of NBS-LRR genes, leading to the expansion of resistance genes [11]. LTR-Rs are abundant in plant genomes including those of Solanaceae crops, such as *Nicotiana sylvestris* (∼38.16%) [12], pepper (more than 70.0%) [13], potato (62.2%) [14], tomato (50.3%) [15] and Petunia (more than 60%) [16]. The role of LTR-Rs in the genome of *S. aethiopicum* is unknown. Whether the resistance of *S. aethiopicum* is raised through LTR-Rs amplification remains to be investigated. Overall, a reference genome for *S. aethiopicum*, as well as for other orphan crops, is greatly needed to advance their research and breeding.

Here we report a draft assembly and annotation for *S. aethiopicum* genome. We found two amplification of LTR-Rs that occurred around 1.25 and 3.5 million years ago resulting in the expansion of resistance genes. We also re-sequenced two *S. aethiopicum* genotypes, “Gilo” and “Shum”, at a high depth (∼60 X) and identified 14,995,740 SNPs, 41,046 of which are closely linked to resistance genes, and subsequently generated a pan-genome of *S. aethopicum*. The genomic data provided in this study will greatly advance the research and breeding activities of the African eggplant.

## Data Description

We sequenced the genome of *S. aethiopicum* using a whole-genome shotgun (WGS) approach. A total of 242.61 Gb raw reads were generated by sequencing the libraries with insert sizes of 250 and 500 bp, and mate-pair libraries with sizes ranging from 2,000 to 20,000 bp on an Illumina Hiseq 2000 platform. The filtered reads that were used for downstream analysis are shown in Supplementary Table S1. The K-mer (K=17) analysis [17] revealed that the *S. aethiopicum* genome is diploid and homozygous with a genome size of 1.17 Gb (Figure S1). We used all the “clean reads” of 127.83 Gb (∼ 109 X) to assemble the genome using Platanus [18] (see methods), and obtained a final assembly of 1.02 Gb in size containing 162,187 scaffolds with N50 values of contigs and scaffolds of 25.2 Kbp and 516.15 Kbp (Table 1 and Supplementary Table S2), respectively. Our results reveal that the *S. aethiopicum* genome is larger than other *Solanum* genomes including tomato (0.76 Gb) and potato (0.73 Gb) [14, 15], while its GC ratio (33.12%) is comparable (Supplementary Table S3).

**Table 1.** The statistics of *S. aethiopicum* genome and gene annotation.

|  |  |
| --- | --- |
| Number of scaffolds | 162,187 |
| Total length of scaffolds | 1.02 Gb |
| N50 of scaffolds | 516.1 Kb |
| Longest scaffolds | 2.94 Mp |
| Number of contigs | 231,821 |
| Total length of contigs | 936 Mb |
| N50 of contigs | 25.2 Kb |
| Longest contigs | 366.2 kb |
| GC content | 33.13% |
| Number of genes | 34,906 |
| Average/total coding sequence length | 1104.3bp/38.5 Mb |
| Average exon/intron length | 265.8bp/613.1 bp |
| Total length of transposable elements | 805.7 Mb (78.23%) |

Repetitive elements, predominantly transposable elements (TE) (Supplementary Table S4), occupied 790 Mbp (76.2%) of the sequenced genome. A majority of the annotated TEs were retrotransposon elements including long terminal repeat elements (LTR), short interspersed elements (SINEs) and long interspersed elements (LINEs). These retrotransposons together occupied 75.42% of the assembly. DNA transposons were also annotated, which accounted for 2.87% of the genome. Among these annotated TEs, LTR-Rs were extraordinarily abundant and occupied 719 Mbp, accounting for approximately 70% of the genome, followed by LINEs and SINEs (Supplementary Table S4).

The protein coding gene models were predicted by a combination of homologous search and *ab initio* prediction. The resulting models were pooled together to generate a final set of 34,906 protein-coding genes. The predicted gene models had an average of 3,038 bp in length, each with an average of 3.15 introns. The average length of coding sequences, exons and introns were 1,104 bp, 265 bp and 613 bp, respectively (Table 1, Supplementary Table S5 and Figure S2). These gene features were, as expected, similar to those in other released genomes including *A. thaliana* [19], and other Solanaceae crops including *S. lycopersicum, S. tuberosum, C. annuum* and *N. sylvestris* [12, 14, 15, 20] (Supplementary Table S5). We further assessed the annotation completeness of this assembly by searching for 1,440 Core Embryophyta Genes (CEGs) with Benchmarking Universal Single-Copy Orthologs (BUSCO, version 3.0) [21]. We found 80.4% CEGs in this assembly with 77.8% being single copies while 2.6% were duplicates (Supplementary Table S6). We also annotated the non-coding genes by homologous search, leading to the identification of 128 microRNA, 960 tRNA, 1,185 rRNA and 503 snRNA genes (Supplementary Table S7).

We annotated a total of 31,863 (91.28%) proteins for their homologous function in several databases. Homologs of 31,099 (89.09%), 26,319 (75.4%), 20,932 (59.97%) proteins were found in TrEMBL, InterPro and SwissProt databases, respectively (Supplementary Table S8). The remaining 3,043 (8.72%) genes encoded putative proteins with unknown functions.

## Analyses

### Genome evolution and phylogenetic analysis

By comparing with other four sequenced Solanaceae genomes of *S. melongena, S. lycopersicum, S. tuberosum* as well as *C. annuum*, 25,751 of the *S. aethiopicum* genes were clustered into 19,310 families using OrthoMCL (version 2.0) [22], with an average of 1.33 genes each. Single-copy genes that were shared by these five genomes were concatenated as a super gene representing each genome and were used to build the phylogenetic tree (Figure 1A). The split time between *S. aethiopicum* and *S. melongena* was estimated to be ∼2.6 million years ago. One hundred and eighty two (182) syntenic blocks were identified by McScanX [23]. We detected evidence of whole genome duplication (WGD) events in this genome by calculating the pairwise synonymous mutation rates and the rate of four-fold degenerative third-codon transversion (4DTV) of 1,686 paralogous genes in these blocks. The 4DTV distribution plot displayed two peaks, around 0.25 and 1, indicating two WGDs (Figure 1B). The first one (peaked at 1) represents the ancient WGD event that is shared by asterids and rosids [24], while the second WGD event is shared by Solanaceae plants suggesting its occurrence predates the split of Solanaceae.

**Figure 1.**
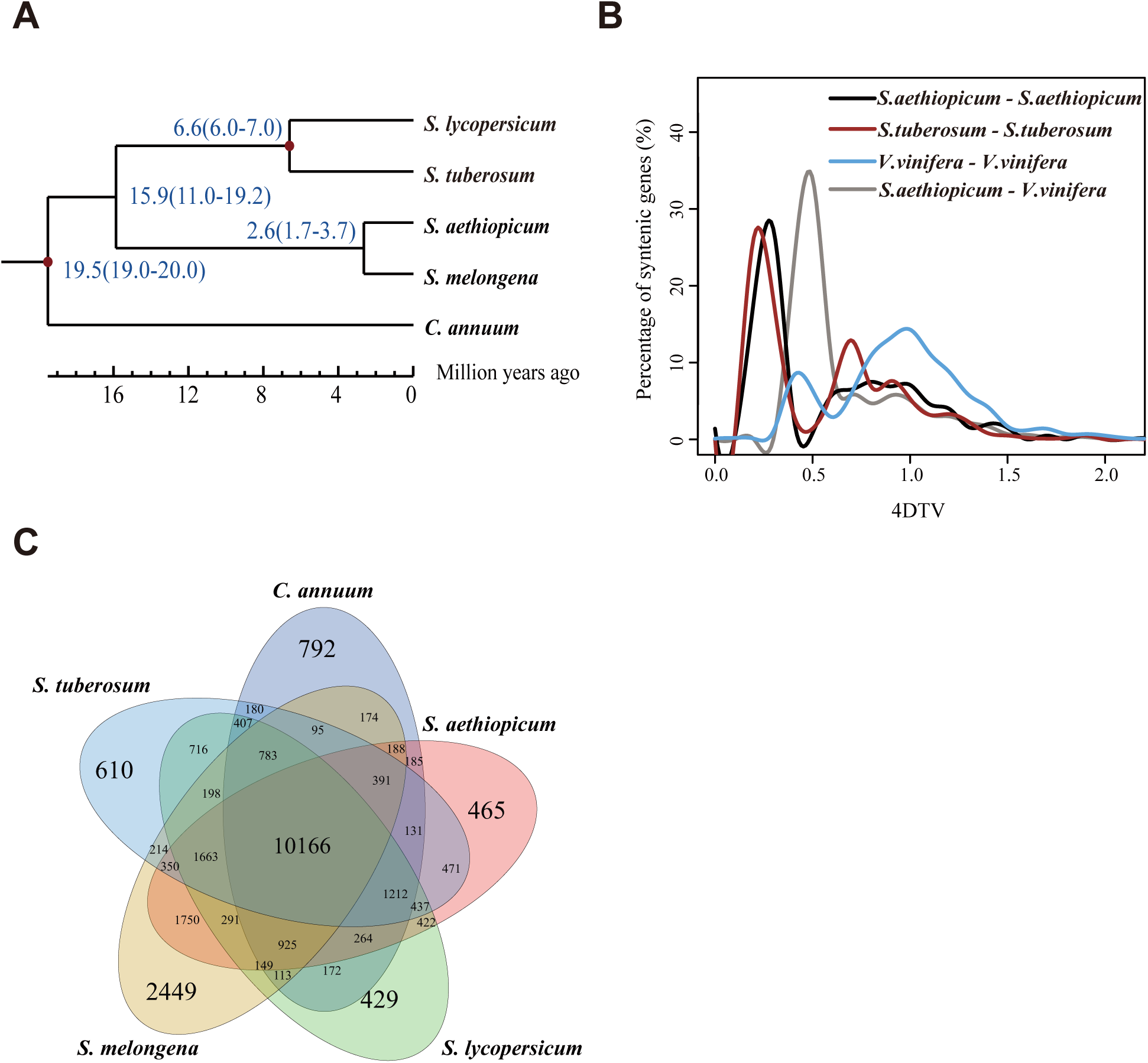
Comparative analysis of the *Solanum aethiopicum* genome. (A) Phylogenetic analysis among *S. melongena, S. lycopersicum, S. tuberosum, S. aethiopicum* and *C. annuum* by using the single-copy gene families. The species differentiation time between *S. aethiopicum* and *S. melongena* was 2.6 MYA (B) Distribution of 4DTv distance, which showed two peaks around 0.25 and 1 (black line), representing two whole genomic duplication events (C) Venn diagram showing overlaps of gene families between *S. melongena, S. lycopersicum, S. tuberosum, S. aethiopicum* and *C. annuum*. A total of 465 gene families were unique to *S. aethiopicum* and 10166 were common shared by the 5 species genome.

### Evolution of Gene families

OrthoMCL [22] clustering of genes from *S. aethiopicum, S. melongena, S. lycopersicum, S. tuberosum* and *C. annuum* identified 25,751 gene families, among which 465 gene families were unique to *S. aethiopicum* and 10,166 were commonly shared (Supplementary Table S9, Figure 1C). As expected, the number of shared gene families decrease as a function of evolutionary distance between *S. aethiopicum* and the selected species (Supplementary Table S10). For example, *S. aethiopicum* shared 15,723 gene families with *S. melongena* as compared to only 13,461 genes shared with *C. annuum*. To further investigate the evolution of gene families, we identified expanded and contracted gene families. Compared to *S. melongena*, 437 gene families were expanded and the majority of the expanded gene families were found to be involved in biological processes related to drought or salinity tolerance as well as disease resistance including defense response (GO:0006952), response to oxidative stress (GO:0006979), glutamate biosynthetic process (GO:0006537) and response to metal ion (GO:0010038) (Supplementary Table S11). On the other hand, there was no gene family contracted when comparison was made with *S. melongena*.

### Amplification of LTR-Rs

LTR retrotransposons (LTR-Rs) comprised ∼70% of the genome and accounted for 89.31% of the total TEs in *S. aethiopicum* (Supplementary Table S4). Consistent with previous studies of LTR-Rs, a majority of the LTR-Rs were classified into *Ty3/Gypsy* (account for 82.36% of total LTR-Rs) and *Ty1/Copia* (account for 14.90% of total LTR-Rs) subfamilies. The proportion of *Ty3/Gypsy* and *Ty1/Copia* LTR-Rs in *S. aethiopicum* is also comparable to those reported in other Solanaceae genomes. To investigate the roles of LTR-Rs in the evolution of *S. aethiopicum*, we detected 36,599 full-length LTR-Rs using LTRharvest [25] with the parameters “-maxlenltr 2000,–similar 75” and LTRdigest software [26]. We further analyzed their evolution, activity and potential biological functions.

The age of each LTR retrotransposon was inferred by comparing the divergence between the 5’ and 3’ LTR-Rs using a substitution rate of 1.3e-8 year^-1^site^-1^ [27]. Two amplifications of LTR retrotransposons were found in *S. aethiopicum* while only one was detected in tomato and hot pepper (Figure 2A). The early amplification occurred at around 3.5 million years ago (MYA), coincident with the LTR-Rs burst found in *C. annuum* [11] (Figure 2A); the second amplification was 1.25 MYA coinciding with the burst in tomato genome [15] (Figure 2A). Although the time of LTR-Rs amplification is vertically coincident between different species, they occurred separately in each genome since *S. aethiopicum* and hot pepper had split about 20 MYA (Figure 1A), and about 4 MYA between *S. aethiopicum* and tomato (Figure 1A). These results imply that environmental stimulators shared between these species during their evolution could have triggered the amplifications observed. We also estimated the amplification time of *Ty3/Gypsy* and *Ty1/Copia* LTR-Rs and found two peaks of around 1.25 and 3.5 MYA for Gypsy LTR-Rs (Figure 2B), while only one peak (around 1.25 MYA) for *Ty1/Copia* LTR-Rs (Figure 2C). Compared with the amplification time of *Ty3/Gypsy* and *Ty1/Copia* LTR-Rs in different species, we found that the insertion time of *Ty1/Copia* LTR-RTs in *S. aethiopicum* and tomato were earlier than that of *S. melongena* and hot pepper. On the contrary, the insertion time of *Ty3/Gypsy* LTR-RTs (around 3.5 MYA) in *S. aethiopicum* was consistent with the insertion time of hot pepper (Figure 2B,2C).

**Figure 2.**
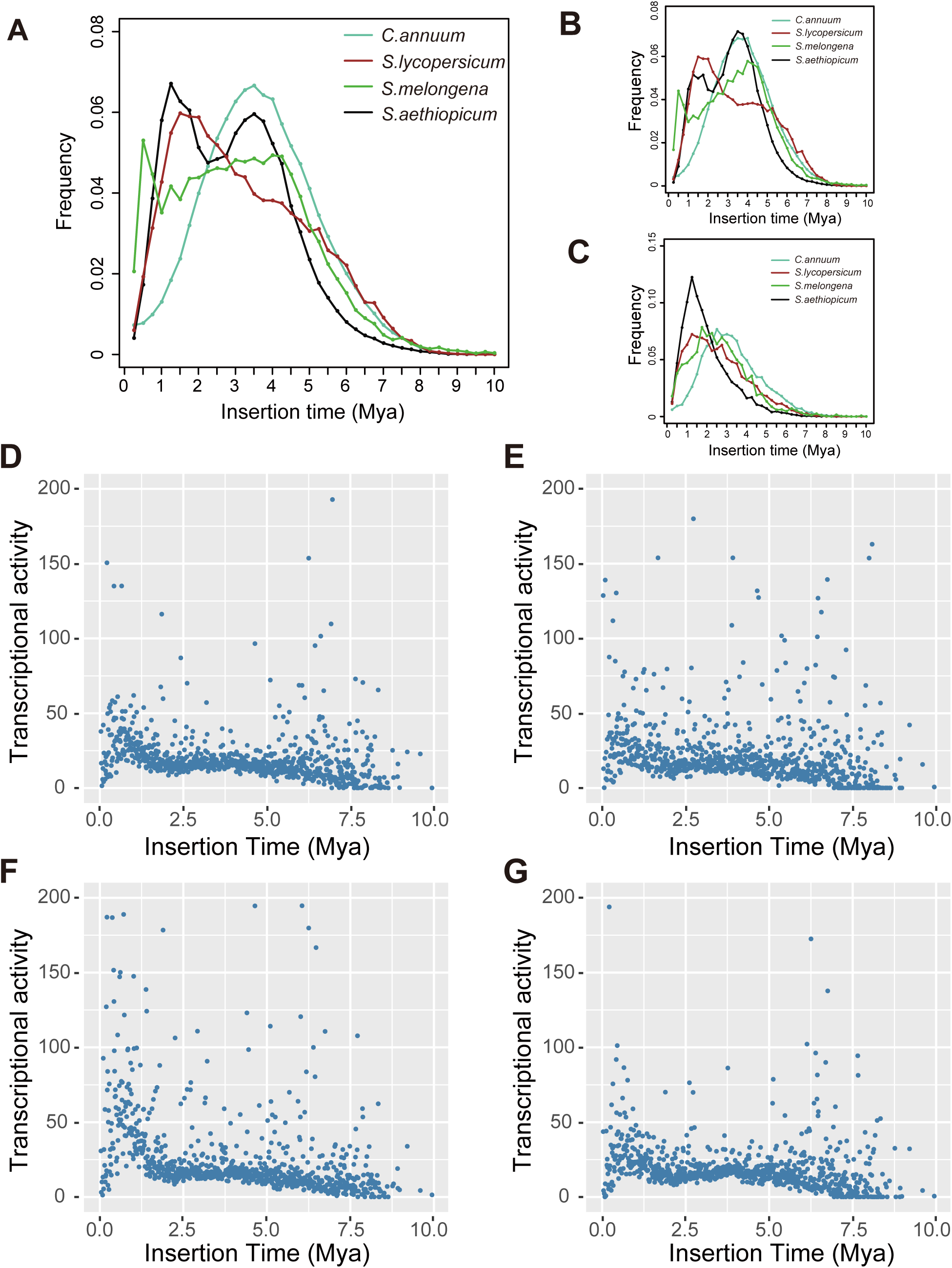
LTR-Rs insertion time distribution and the expression level of LTR-Rs in different tissues. Insertion time distribution of total LTR-Rs (A), *Ty3/Gypsy* LTR-Rs (B) and *Ty1/Copia* LTR-Rs (C) of *C. annuum, S. melongena, S. lycopersicum* and *S. aethiopicum*. The x- and y-axes respectively indicate the insertion time and the frequency of inserted LTR-Rs. The expression levels of LTR-Rs in Flower (D), Fruit (E), Leaf (F), Root (G).

To investigate the activities of these LTR-Rs, we measured their expression levels by using RNA-seq data from different tissues (see method). The younger LTR-Rs were expressed in higher levels than those of older LTR-Rs. We detected two peaks of LTR-Rs activities at positions corresponding to the two rounds of LTR-Rs insertions (Figure 2D-G). The slight shift of the former peaks indicates that the activities degenerated slower than the sequences of LTR-Rs (Figure 2D-G). The LTR-R activities varied across these tissues. The degeneration of LTR-R activities was slower in fruits and roots, compared to those in flowers and leaves (Figure 2D). This pattern was also confirmed by the varied activity of each LTR-Rs across these tissues (Figure 2D), which implies that these LTR-Rs play different roles in development.

### Increased resistance is facilitated by LTR-Rs amplification

We identified 1,156 LTR-Rs captured genes and 491 LTR-Rs disrupted genes. The insertion time of LTR-Rs captured and LTR-Rs disrupted genes both ranged from 1.5 to 3.5 MYA (Figure 3A), showing a pattern similar to the whole LTR-Rs insertions (Figure 2A). These results suggest that gene disruption and capturing mediated by LTR-Rs occurred simultaneously. We further classified the LTR-Rs captured genes into gene ontology (GO) categories and performed GO enrichment analysis. GO terms related to disease resistance including “defense response to fungus (GO:0006952)”, “chitin catabolic process (GO:0006032)”, “chitinase activity (GO:0004568)”, “chitin binding (GO:0008061)”, “cell wall macromolecule catabolic process (GO:0016998)” and “defense response to bacterium (GO:0042742)” were significantly overrepresented in the LTR-Rs captured genes (Figure 3B, Supplementary Table S12), suggesting a likely role of enhancing disease resistance.

**Figure 3.**
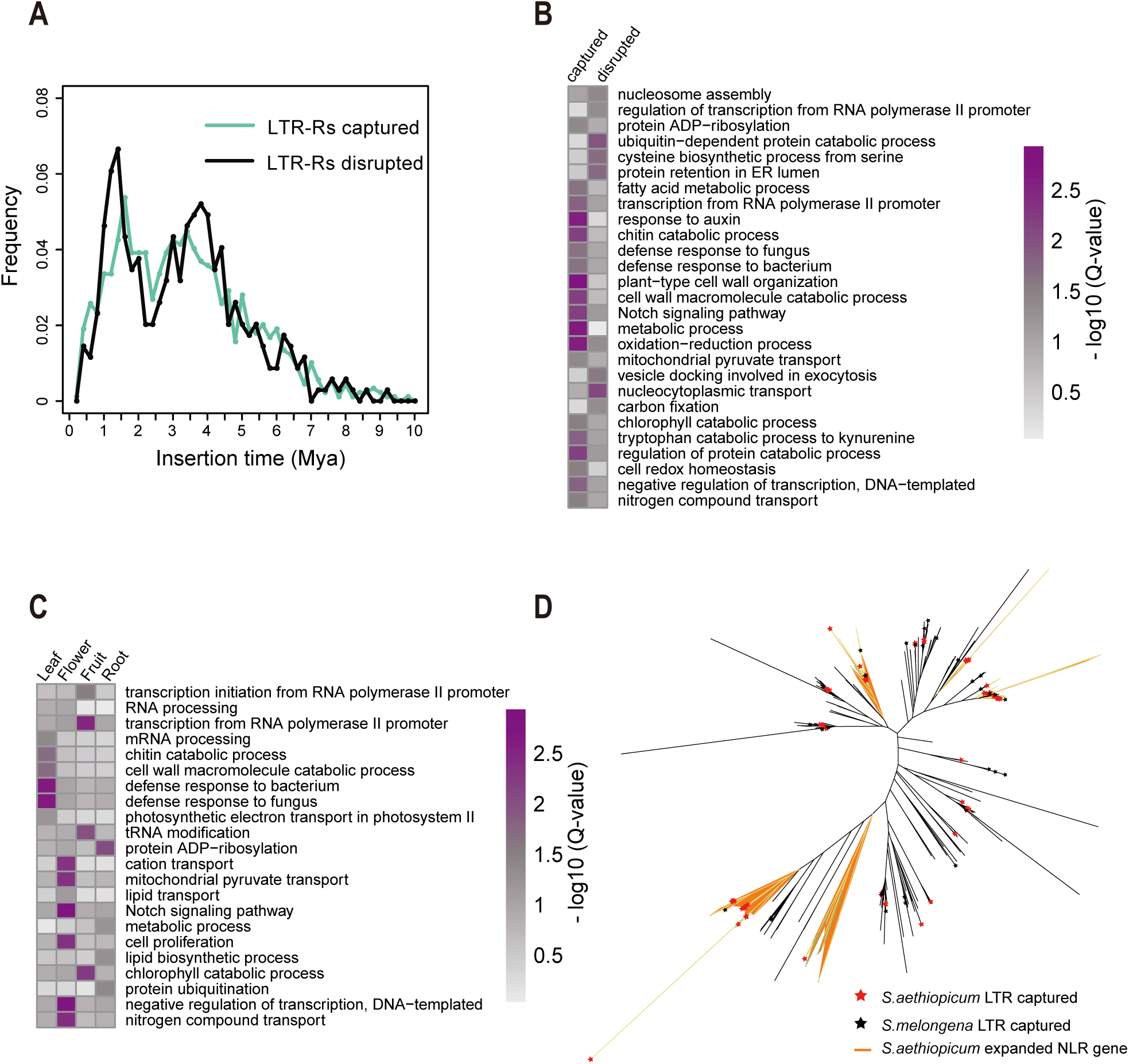
LTR-Rs captured and disrupted genes. (A) The distribution of ages of LTR-Rs captured and disrupted genes. (B) GO enrichment analysis between the LTR-Rs captured and disrupted gene set. (C) GO terms enriched in LTR-Rs captured genes that are specifically highly expressed in various tissues including leaf, flower, root and fruit. (D) Phylogenetic tree of NLR gene in *S. aethiopicum* and *S. melongena*.

We also analyzed the expression of genes captured by LTR-Rs. It was intriguing to find that a majority of these genes were specifically active in only one tissue (Figure S3). Among these genes, 159 (13.75%), 105 (9.08%), 106 (9.16%) and 129 (11.15%) were specifically highly expressed in root, leaf, flower and fruit, respectively. We observed that the genes captured by LTR-Rs that were specifically active in leaf tissues were significantly enriched in functions related to disease resistance (Supplementary Table S13). The biological processes and molecular activities related to disease resistance mentioned above were overrepresented in these genes (Figure 3C). The high expression level of resistance genes in leaves would arm the plant with stronger resistance to pathogens. On the contrary, these GO terms were not enriched in the total leaf-specifically highly expressed genes. Instead, as expected, “photosynthesis” and “photosystem I” were significantly overrepresented (Supplementary Table S14). The discrepancy between these two gene sets highlights the contribution to resistance of LTR-captured genes.

Proteins containing nucleotide-binding site and leucine-rich repeat domains (NBS-LRRs) are major components responsible for defense against various phytopathogens [28]. The NBS-LRRs family is highly expanded in plants with numbers ranging from less than 100 to more than 1000 [29, 30]. As NBS-LRR genes are often co-localized with LTR-Rs [31], we inspected their genomic locations in *S. aethiopicum* genome. We identified a total of 447 NBS-LRR genes in the genome, among which 62 (13.8%) NLR genes co-localized with LTR-Rs were identified as LTR-Rs captured genes. A similar percentage (∼13%) of LTR-captured NLR genes was also reported in hot pepper [11]. The phylogenetic tree shows a substantial expansion of NLRs after the amplification of LTR in *S. aethiopicum* (Figure 3D). A similar expansion was also observed in eggplant. However, the number was significantly fewer than that in *S. aethiopicum*, probably due to the limited number of LTR-Rs in eggplant genome (Supplementary Table S15).

### Polymorphisms in different *S. aethiopicum* groups

We totally sequenced 60 *S. aethiopicum* genotypes in two major groups, “Gilo” and “Shum” and 5 accessions of *S. anguivi*, the ancestor of *S. aethiopicum*. By sequencing each accession with ∼ 60 Gb raw data (60 X) (Supplementary Table S16), we identified a total of 18,614,838 SNPs and 1,999,241 indels, with an average of 3,530,488 SNPs for each accession. On average, there were 18,090 SNPs and 1,943 indels per megabase. Among them, 426,401 (2.07%), 821,101 (3.98%) and 19,374,353 (93.99%) were located in exons, introns and intergenic regions respectively (Table 2). There were 267,710 SNPs that resulted in changes of amino acid sequences by introducing new start codons, premature stop codons, or nonsynonymous substitutions (Table 2). In addition, we also identified 1,999,241 indels and 1,255,302 structural variations (SVs). Of the detected indels, 178,260 (8.90%) were located in genic regions, among which 2,977 (0.13%) caused frame shift changes and, therefore, resulted in changes of amino acid sequences that may have led to malfunction of genes. Furthermore, 106,377 SVs were identified in gene regions including 53,736 (50,51%) deletions, 34,368 (32,31%) insertions and 8,872 (8.34%) duplications.

**Table 2.** Statistics of SNPs and indels for 65 accession

|  |  | Number | Percentage (%) |
| --- | --- | --- | --- |
| SNPs | Exon | 393,882 | 2.12 |
|  | Intron | 675,360 | 3.63 |
|  | Intergenic | 17,552,823 | 94.29 |
|  | Synonymous | 126,172 | 0.67 |
|  | Non-synonymous | 267,710 | 1.44 |
|  | Total | 18,614,838 |  |
| Indels | Exon | 32,519 | 1.62 |
|  | Intron | 145,741 | 7.28 |
|  | Intergenic | 1,821,530 | 91.11 |
|  | Frame shift | 2,977 | 0.13 |
|  | Total | 1,999,241 |  |

We counted the SNPs and indels in each group. As a result, 12,777,811, 15,165,053 and 8,557,818 SNPs were found in “Gilo”, “Shum” and “*S. anguivi*”, accounting for 68.64%, 81.47% and 45.97% of the total SNPs, respectively. There were, 2,019,539 (10.85%%), 4,747,418 (25.50%) and 587,885 (3.16%) SNPs that were unique to “Gilo”, “Shum” and “*S. anguivi*” respectively (Figure 4A). The majority (93.13%) of SNPs in “*S. anguivi*” were shared with either “Gilo” or “Shum” (Figure 4A), which is in line with the fact that “*S. anguivi*” is the ancestor [32]. Similarly, 92.62% of the indels identified in “*S. anguivi*” were also share with “Gilo” or “Shum” (Figure 4B).

**Figure 4.**
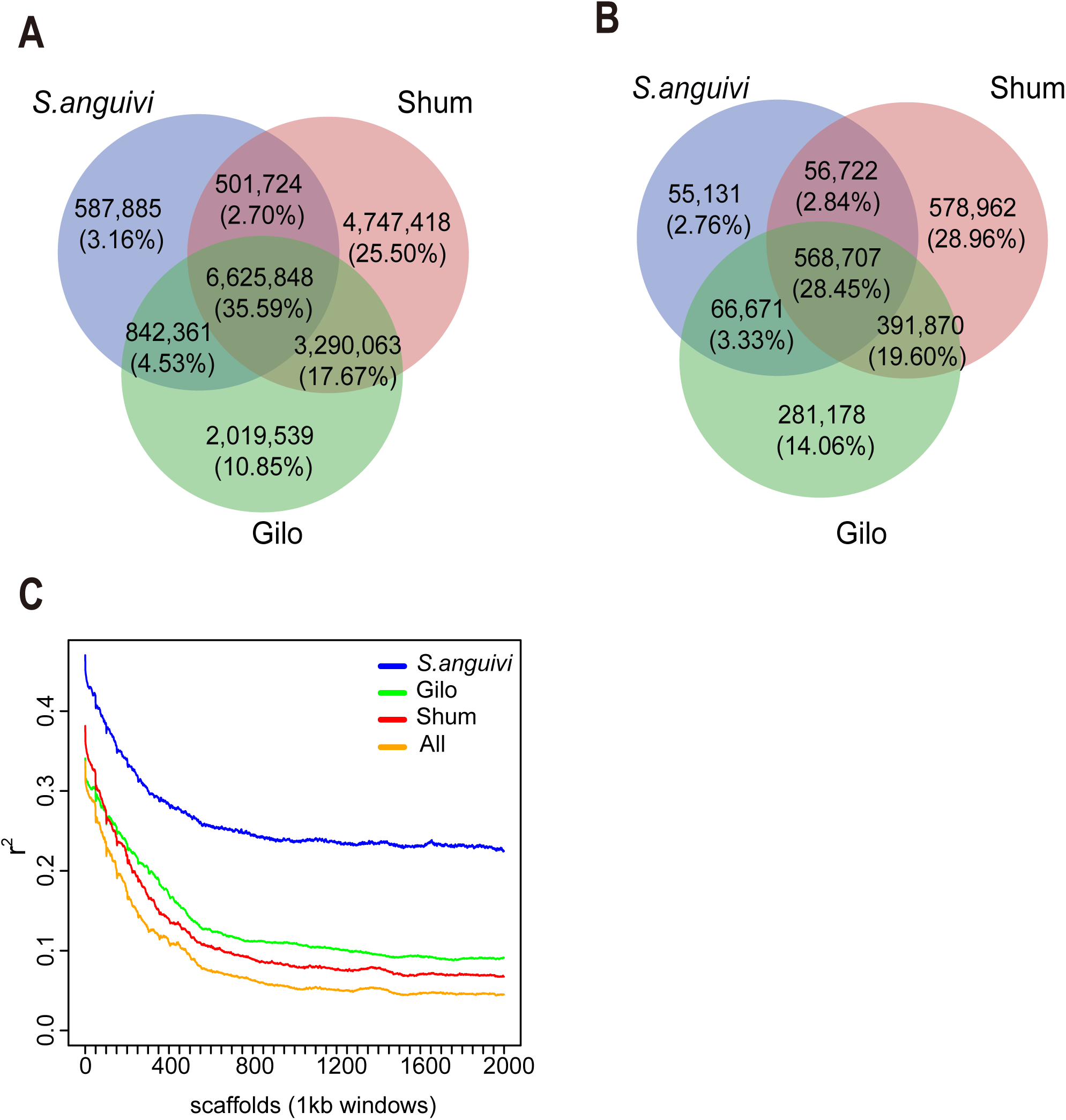
SNPs, Indel and LD decay for “Gilo”, “Shum” and “*S. anguivi*” groups. (A) 2,019,539 (10.85%%), 4,747,418 (25.50%) and 587,885 (3.16%) SNPs that were unique to “Gilo”, “Shum” and “*S. anguivi*” respectively and the majority (93.13%) of SNPs in “*S. anguivi*” were shared with either “Gilo” or “Shum”; (B) 14.06%, 28.96% and 2.76% indels were unique to “Gilo”, “Shum” and “*S. anguivi*” respectively and similar to the statistics of SNPs in these groups, 92.62% of indels in “*S. anguivi*” were shared with either “Gilo” or “Shum”; (C) Linkage disequilibrium (LD) estimation revealed that r^2^ reaches the half maximum value at ∼150 kb.

Nucleotide diversity (π) of all the genotypes was determined to be 3.58 × 10^-3^ for whole genomes, 2.06 × 10^-3^ for genic regions and 3.75 × 10^-3^ for intergenic regions. Nucleotide diversity for each genotype revealed lower diversity for “Gilo” (*S. anguivi*: 3.16 × 10^-3^, Shum: 3.65 × 10^-3^ and Gilo: 2.55 × 10^-3^). Linkage disequilibrium (LD) estimation using Haploview (version 4.2) [33] revealed that r^2^ reaches the half maximum value at ∼150 kb (Figure 4C), which is smaller than in other Solanaceae crops, for example, tomato (2,000 kb) [34]. Since *S. aethiopicum* has been routinely used to improve disease resistance in eggplant and other Solanaceaee crops [10], we further identified SNPs strongly associated with resistance genes by selecting those within 150 kb of resistance genes. A total of 5,562 SNPs were finally selected (205 genes), which could be used to assist the selection of plants with disease resistance (Supplementary Table S16).

### Population structure and demography of *S. aethiopicum*

To investigate the evolution and population demography of *S. aethiopicum*, we first built a maximum-likelihood (Figure 5A) phylogenetic tree using the full-set of SNPs. We observed population structure in the genome-wide diversity. As anticipated, the accessions from “Gilo” and “Shum” were clearly separated in the tree, with only one exception in each group, probably due to labelling errors. On the other hand, accessions of “*S. anguivi*”, which is the known ancestor of *S. aethiopicum*, did not cluster separately, but instead, grouped with either “Gilo” or “Shum”. This structure was also supported by principal-component analysis (PCA), which clearly separated these accessions into two clusters (Figure 5B and Figure S4).

**Figure 5.**
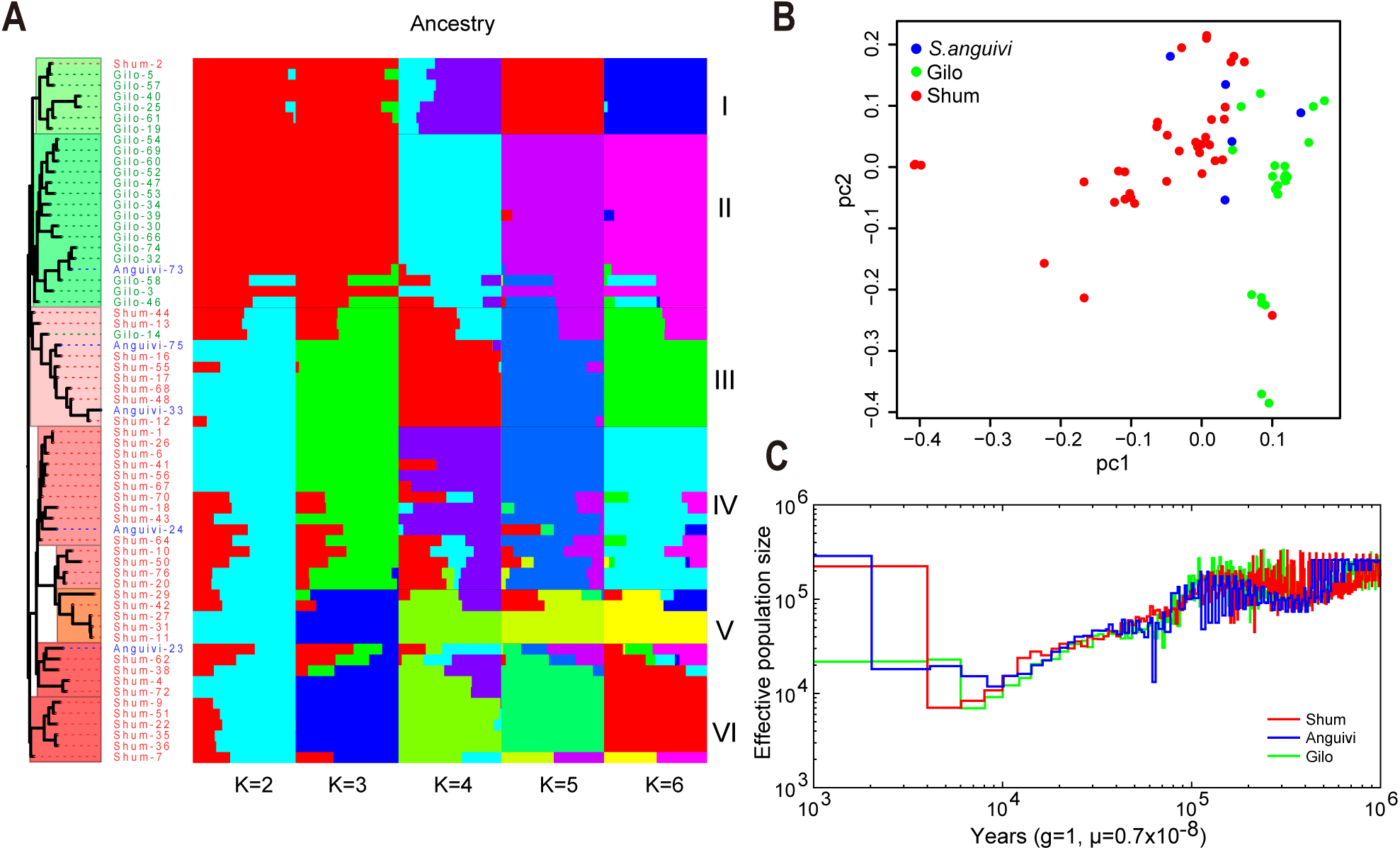
Population structure and demography of *S. aethiopicum*. (A) A maximum-likelihood phylogenetic tree using the full-set of SNPs; (B) Principal-component analysis (PCA); (C) PSMC analysis indicated a distinct demographic history of *S. aethiopicum* from 10,000 to 100 years ago, in which a bottleneck was shown around 4,000-5,000 years ago, followed by an immediate expansion of population size.

The domestication history of *S. aethiopicum* was also inferred by constructing a multi-level population structure using ADMIXTURE [35], which enabled us to estimate the maximum likelihood ancestry (Figure 5A). The parameter K, representing the number of sub groups to be divided, were set from 2 to 9, and the cross-validation error was calculated individually. The cross-validation (CV) error converged to 0.4375 when K = 6, therefore, the population was divided into six sub-groups, I, II, III, IV, V and VI (Figure 5A). The structure changes along with the increase of K from 2 to 6 showed a time lapse domestication history of *S. aethiopicum*, which was first split into two groups, “Gilo” and “Shum”. The former was subsequently divided into sub-groups I and II. Two groups emerged in “Shum” when K = 3, each of which was then divided into two sub-groups when K = 6. In summary, “Gilo” was divided into two sub-groups (I and II) and “Shum” was divided into four sub-groups (III, IV, V and VI).

Furthermore, the demographic history of *S. aethiopicum* was inferred using the pairwise sequential Markovian coalescent model PSMC [36]. By doing this, we inferred the changes of effective population sizes of *S. aethiopicum* (Figure 5C). Our data revealed the distinct demographic trends from 10,000 to 100 years ago, in which a bottleneck was shown around 4,000-5,000 years ago, followed by an immediate expansion of population size. The great expansion of population might be due to the early consumption of *S. aethiopicum* in Africa because it is coincidence with the human population growth in western Africa occurring 4,000-5,000 years ago [37].

### Artificially selected genes in *S. aethiopicum*

We used *ROD* and *Fst* measure to detect artificially selected regions along the genome. Briefly, *ROD* and *Fst* were calculated in a sliding non-overlap 10 kb-window, and regions with *ROD* > 0.75 and *Fst* > 0.15 were identified as candidate regions under selection. As a result, genomic regions of 3,238 and 1,062 windows were found to be under selection during the domestication of “Gilo” and “Shum”, respectively (Supplementary Table S17). Among them, 161 windows were commonly shared in these two groups while 3,077 and 901 windows were unique to “Gilo” and “Shum”, respectively. Genes located at these regions were identified as selected genes. A number of 1,406 and 36 selected genes were identified in “Gilo” and “Shum”, respectively, of which 12 genes were selected in both. GO enrichment analysis showed that genes selected in both the groups were enriched in “transport” (Supplementary Table S18). In addition, GO terms of “response to auxin”, “response to hormone”, “response to salt stress” and “response to water” were overrepresented in genes selected only in “Gilo” or “Shum”. This result explains the enhanced tolerance to drought and salinity in *S. aethiopicum*.

Furthermore, we focused on the diversity of genes co-localized with LTR-Rs. A number of 24,682 SNPs were located within these genes, corresponding to 0.133% of the total number of SNPs (18,614,838), which is substantially fewer than would be expected if SNPs were evenly distributed in all the genes, especially because the LTR-R co-localized genes comprise 3.31% of the total gene set. The repellant of SNPs in these genes suggests purifying selection, which was also supported by the large amount (9,728, 39.41%) of rare SNPs (minor allele frequency < 5%) found in these genes. Nevertheless, we also noticed that nonsynonymous SNPs (9,544) were much more abundant than synonymous ones (5,310) in these genes. These variations led to amino acid changes in the encoded proteins, which may have contributed to the diversification of resistance genes.

### Pan- and core-genome of *S. aethiopicum*

Gene content varies across different accessions. A single reference assembly is not sufficient to include all the genes of *S. aethiopicum*. Therefore, we assembled contigs for each individual accession using pair-ended reads with coverages of 30 - 60 X (Supplementary Table S19).

We assembled the genomes individually using SOAPdenovo2 [38] and filtered out contigs smaller than 2 kb. As a result, 753,084 contigs were retained, among which 432,785 were from “Shum”, 260,119 were “Gilo” and 60,180 were from “*S. anguivi*”. These contigs were further pooled separately and cleaned by removing duplicates using CD-HIT [39], which led to the retention of 97,429, 76,638 and 36,915 contigs for “Shum”, “Gilo” and “*S. anguivi*”, respectively. The annotation of these contigs resulted in 41,626, 22,942 and 17,726 protein-coding genes, among which we identified accessory gene sets of 29,389, 23,726 and 12,829 for “Shum”, “Gilo” and “*S. anguivi*”, respectively, by comparing with the reference genome. We generated a pan-genome of *S. aethiopicum* of 51,351 genes (Supplementary Table S20). The average length of accessory genes was 1.62 kb with 2.22 introns, comparable to gene models in the reference, suggesting they were accurately annotated. We further annotated their putative functions by querying against protein databases. As a result, a total of 48,572 (94.59%) genes were annotated with function descriptions (Table 20 and Supplementary Table S21). Among all the identified gene models, 21,711 (44.28%) were commonly shared by these three groups and defined as core genes. As expected, they were majorly composed of house-keeping genes (Supplementary Table S22). However, a caveat should be noted that the number of core genes was under estimated because “*S. anguivi*” was inadequately represented and the other two groups of *S. aethiopicum*, “Kumba” and “Aculeatum”, were not included in this study.

## Discussion

As a close relative, *S. aethiopicum* is compatible to *S. melongena* and routinely used to improve disease resistance of *S. melongena* by providing resistance genes [10]. The genomic analysis of *S. aethiopicum* revealed expansion of resistance gene families compared to its close relatives including tomato, potato, eggplant and hot pepper, which is mediated by LTR amplifications. LTR amplification is one of the major forces driving genome evolution. It shapes the genome by capturing, interrupting or flanking genes [40]. The consequences of LTR insertions depend on the genomic position of insertion. For example, inserting into protein coding sequences results into the pseudogenization. LTR-Rs adjacent to protein coding genes can down regulate or silence the expression of flanking genes by extending methylation region or by producing anti-sense transcripts [41-44]. In addition, LTR-Rs also mediate gene retroposition, capturing genes back into the genome [40]. During LTR amplification, LTR preferentially captured genes related to disease resistance. GO terms of related to disease resistance were over-represented in LTR-captured genes. Particularly, the enrichment of terms of “chitin binding (GO:0008061)” and “chitinase activity (GO:0006032)” (Figure 3B, Supplementary Table S12) implies that these genes were selected to resist the infection of fungal pathogens, such as *Fusarium oxysporum* [45]. On the contrary, there was no GO term enriched in genes disrupted by LTR-Rs. It seems gene disruption by LTR-Rs is a random event in term of gene function. The age distribution of LTR-Rs captured genes coincidently fit with that of the LTR-disrupted genes, suggesting they were occurred simultaneously (Figure 3A). Why genes related to resistance were favored by LTR-Rs? An explanation to this bias is that these genes were more active than other genes when LTR retrotransposition occurred. The expression pattern of LTR-Rs captured genes varied among different tissues. Those related resistance are specifically active in leaf, while those engaged in transport of cation, nitrogen transport and cell proliferation are active in flower. These data suggest low abundance of transcripts of resistance genes, therefore small chance to be captured, in flower under normal conditions. A possible scenario is that LTR retrotransposition occurred under stressed conditions, which activated the expression resistance genes in gamete, and simultaneously the activity of LTR retrotransposition. These stresses could be extreme temperature or pathogen infection and so on. In another organism, a “reinforcement model” was proposed to explain the accumulation of stress responsive genes in the genome, in which environmental stress activates both the expression of stress responsive genes and the activities of retrotransposons, resulting in higher probabilities of retrotransposition for stress responsive genes [46, 47].

There were four major groups of *S. aethiopicum*, “Gilo”, “Shum”, “Kumba” and “Aculeatum”. In this work, we re-sequenced accessions from the former two groups, which were widely consumed. The accessions re-sequenced in this study was divided into six sub-groups (2 for “Shum” and 4 for “Gilo”). By scanning for regions with lower diversities in the genome, we identified regions and genes under selection during domestication of *S. aethiopicum*. A number of genes involved in responses to salt, water and drought tolerance were selected. Furthermore, purification selection was also found in resistance genes.

In this work, the genomes of *S. aethiopicum* accessions were sequenced with high depth (30 – 60 X) (Table S19), which enabled us to assemble the contigs for each individual. Despite there were only genotypes from two groups were included in this study, we intended to supplement the reference gene set with accessory genes by pooling these contigs together for gene prediction and annotation. These “pan-genome” would provide a more comprehensive understanding of *S. aethiopicum*.

Overall, we present a reference genome for African eggplant, which will provide the basic data resource for the further genomic research and breeding activities within *S. aethiopicum*. For example, the gene sequences annotated in the genome are essential for developing genome editing vectors aiming to create null or weakened mutants. Molecular markers developed using the genome sequences will also enable fast and precise selection of superior accessions by breeders.

### Potential implications

A great number of indigenous crops grown and used locally by smallholders in Africa had played critical roles in alleviating malnutritional and food shortage problems in rural areas of Africa. Most of these crops are hardly known by scientists and consumers out of these areas, and therefore obtained very little investment in scientific research and production. These crops are called “orphan crop”, “neglected crops” or “forgotten crops”. African orphan crop consortium (AOCC) was established in 2012 with an aim to address the food and nutritional security in Africa by using 101 of selected orphan crops that were originated or naturalized in Africa. One effective approach of boosting the research and breeding level of these orphan crops is to sequence their genomes, which would enable genome-assisted selection and precise edition of their genomes. Five genomes of orphan crops including *Vigna subterranea, Lablab purpureus, Faidherbia albida, Sclerocarya birrea*, and *Moringa oleifera* were recently released [48]. *S. aethiopicum* is also one of these selected orphan crops. The genome sequence released in this work will benefit the research and production of this crop.

## Methods

### DNA extraction, library construction and sequencing, and genome assembly

High molecular genomic DNA was extracted from young leaves of *S. aethiopicum* and then fragmented and used to construct paired-end libraries with insert sizes of 250 bp, 500 bp, 2 kb, 6 kb, 10 kb and 20 kb following standard Illumina protocols. These libraries were then sequenced on an Illlumina HiSeq 2000 platform. A total of 242.61 Gb raw reads were generated by sequencing these libraries. Filtering of duplicated, low quality reads and reads with adaptors was done using SOAPfilter (version 2.2, an application included in the SOAPdenovo package) [38]. We used 17 k-mer counts [17] of high-quality reads from small insert libraries to evaluate the genome size and heterozygosity using GCE [49] and Kmergenie [50]. We assembled the genome using Platanus [18].

Genomic DNA used for re-sequencing was extracted from young leaves of 65 accessions. The DNA was sheared into small fragments of ∼ 200 bp and used to construct paired-end libraries following standard BGI protocols and subsequently sequenced on a BGI-500 sequencer. Ultra-deep data was produced for each accession with coverage ranging from ∼45 to ∼75 (Supplementary Table S19).

### RNA extraction, library construction and sequencing

For RNA extraction, seeds of Gilo and Shum inbred lines were obtained from Uganda Christian University. The seeds were planted in a screen house at the BecA-ILRI Hub (Nairobi, Kenya) in PVC pots (13cm height and 11.5cm diameter) containing sterile forest soil and farmyard manure (2:1). The seedlings were later transplanted into larger PVC pots of 21cm height and 14cm diameter. Plants were raised in a screen house at 21-23°C and 11-13°C day and night temperatures respectively (average 12 light hours per day). The plants were regularly watered to maintain moisture at required capacity.

Two plants were selected randomly from each of Gilo and Shum accessions and tagged at seedling stage for tissue sampling. Fresh tissues were sampled from each of the tagged plants and flash-frozen in liquid nitrogen immediately. Total RNA was extracted from the frozen tissues using the ZR Plant RNA MiniprepTM Kit (Zymo Research, CA, USA) according to the manufacturer’s instructions. RNA integrity was evaluated by electrophoresis in denaturing agarose gel (1% agarose, 5% formamide, 1X TAE) stained with 3x Gel Red (Biotium Inc., CA, USA). The RNA was quantified using the Qubit RNA Assay Kit (Life Technologies, Thermo Fisher Scientific Inc.). Ribosomal RNA (rRNA) was removed from 4µl of total RNA of each sample using Epicentre Ribo-zero(tm) rRNA Removal Kit (Epicentre, Madison, WI, USA). The rRNA-depleted RNA was then used to generate strand-specific RNA-seq libraries using TruSeq^®^ Stranded mRNA Kit (Illumina, San Diego, CA, USA). In total, 20 mRNA libraries were prepared, multiplexed (10 samples at a time) and sequenced as paired-end reads on the MiSeq (Illumina) platform at the BecA-ILRI hub.

### Repeat annotation

Tandem repeats were searched in the genome using Tandem Repeats Finder (TRF for short, version 4.04) [51]. Transposable elements (TEs) were identified by a combination of homology-based and *de novo* approaches. Briefly, the assembly was aligned to known repeats database (Repbase16.02) using RepeatMasker and RepeatProteinMask (version 3.2.9) [52] at both the DNA and protein level. In *de novo* approach, RepeatModeler (version 1.1.0.4) [53] was employed to build a *de novo* repeat library using S. aethiopicum assembly, in which redundancies were filtered out before TEs in the genome were identified by RepeatMasker [52]. Long terminal repeats (LTR) were identified using LTRharvest [25] with a criterion of 75% similarity on both sides. LTRdigest [26] was used to identify the internal elements of LTR-Rs with the eukaryotic tRNA library (http://gtrnadb.ucsc.edu/). Identified LTR-Rs includes intact PPT (poly purine tract) and PBS (primer binding site) with long terminal repeats regions (LTR-Rs) on both sides were considered as the final intact LTR-Rs, and were then classified into super-families, *Gypsy* and *Copia*, by querying against Repbase16.02 [54].

### Annotation of gene models and ncRNA

Gene models were predicted with a combination of *de novo* prediction, homology search and RNA-aided annotation. Augustus software [55] was used to perform *de novo* prediction after the annotated repeats were masked in the assembly. To search for homologous sequences, protein sequences of other four closely related species (*S. lycopersicum, S. tuberosum, C. annuum, N. sylvestris*) together with *Arabidopsis thaliana* were used as query sequences to search the reference genome using TBLASTN [56] with the e-value <= 1e-5. Regions mapped by these query sequences were subjected to GeneWise [57] together with their flanking sequences (1000 bp) to identify the positions of start/stop codon and splicing. For RNA-aided annotation, the RNA-seq data from different tissues of *S. aethiopicum* were mapped to the assembly of *S. aethiopicum* genome using HISAT [58]. Good quality transcripts were assembled using StringTie [59]. GLEAN software [60] was used to integrate mapped transcripts from different sources to produce a consensus gene set. tRNAscan-SE [61] was performed to search for reliable tRNA positions. snRNA and miRNA were detected by searching the reference sequence against Rfam database [62] using the Blast software package [56]. rRNAs were detected by aligning with BLASTN [56] against known plant rRNA sequences (www.plantrdnadatabase.com). For functional annotation, protein sequences were searched against the Swissprot, TrEMBL, KEGG (Release 88.2), InterPro, Gene Ontology, COG and Non-redundant protein NCBI databases [63-67].

### Gene family analysis

Proteins of *S. lycopersicum, S. aethiopicum, S. tuberosum, C. annuum* and *S. melongena* were selected to perform all-against-all comparison using BLASTP [56] with e-value cutoff of <=1e-5. OrthoMCL [22] and default MCL inflation parameter of 1.5 was used to define the gene families. Single-copy families were selected to perform multiple sequence alignment by using MAFFT [68]. Fourfold degenerate sites were picked and used to construct a phylogenetic tree based on the maximum likelihood method by PhyML [69] with *C. annuum* as the outgroup. WGD analysis was achieved through the identification of collinearity blocks by paralog gene pairs in MCscanX [23]. Each aligned paralog gene pairs were concatenated to a super-sequence in one collinearity block and 4dTv (transversion of fourfold degenerate site) values of each block were calculated. We also determined the distribution of 4DTv values in order to estimate the speciation between species or WGD events. The divergence time of *S. aethiopicum* was estimated using MCMCtree programme [70] with the constructed phylogenetic trees and the divergence time of *C. annuum* [20] and *S. tuberosum* [14].

### Analysis of LTR-Rs

The insertion times of identified intact LTR-Rs were estimated based on the sequence divergence between the 5’ and 3’ LTR of each element. The nucleotide distance K between one pair of LTR-Rs was calculated using the Kimura 2-parameter method in the Distmat (EMBOSS package) [71]. An average base substitution rate of 1.3e-8 [27] was used to estimate the insertion time based on the formula T = K / 2r [11].

Transcriptomic data were used to analyze the activity of intact LTR-Rs. After filtering and removing the low-quality reads, we mapped the high quality reads from each against the full length LTR-Rs sequence using BWA-MEM software [72] with default parameters. The expression levels of intact LTR-Rs was calculated using EdgeR package [73] and visually presented by the pheatmap in R package [74].

### Analysis of NLR genes

To identify NB-ARC genes in the *S. aethiopicum* genome, we used the HMM profile of the NB-ARC domain (PF00931) as a query to perform an HMMER search (version 3.2.1, http://hmmer.org/) against protein sequences of tomato, potato, hot pepper [14, 15, 20] and annotated sequences of *S. aethiopicum* with e-value cut-off of <=1e-60. The aligned NB-ARC domain sequences of *S. aethiopicum* were extracted and used to build the *S. aethiopicum*-specific HMM model. The NB-ARC domain sequences of tomato, potato and hot pepper were mapped as the query sequences against *S. aethiopicum* genome using TBLASTN [56] with e-value cut-off of <=1e-4 in GeneWise software [57] in order to identify candidate NLR genes at a whole genome level. Final NLR genes were confirmed by searching the genome with *S. aethiopicum*-specific HMM model constructed above with e-value cut-off of <=1e-4. The retroduplicated NLRs were identified according to the method of Kim et al. (2017) and the phylogenetic tree of both *S. aethiopicum* and *S. melongena* NLRs was constructed using FastTree [75] with default parameters.

### SNP Calling

Genome Analysis Toolkit (GATK) pipeline (https://software.broadinstitute.org/gatk/) was used to call SNPs and indels. Briefly, low quality, duplicated and adaptor-contaminated reads were filtered off using SOAPfilter (version 2.2) [38] before further processing. To reduce the computing time, we sequentially linked scaffolds in the assembly into 24 pseudo-chromosomes, in which the original scaffolds were separated by 100 Ns before mapping of reads using BWA [72] with default parameters. Picard-tools (https://broadinstitute.github.io/picard/) and SAMtools [76] were used to further process the alignment outputs including sorting and marking of duplicates. After alignment and sorting, GATK pipeline (version 4.0.11.0, https://software.broadinstitute.org/gatk/) was used to call SNPs by sequentially implementing the following modules: RealignerTargetCreator, IndelRealigner, UnifiedGenotyper, samtools mpileup, samtools mpileup, VariantFiltration, VariantFiltration, BaseRecalibrator, AnalyzeCovariates, PrintReads, HaplotypeCaller. This pipeline produced a file in gvcf format displaying the called SNPs and indels, which were further filtered according to genotype information before they were further analyzed using PLINK software [77] for quality control with “GENO>0.05, MAF<0.1, HWE test p-value <=0.0001” parameters. The loci of these SNPs and indels were anchored back to the original scaffolds and annotated using snpEff [78]. To identify structural variations (SVs), sample information was added using AddOrReplaceReadGroups, a module of Picard-tools, and SVs were detected using DiscoverVariantsFromContigAlignmentsSAMSpark, a module of GATK.

### Population analysis

A Maximum-Likelihood phylogenetic tree was constructed based on the genotypes at all the SNP loci using FastTree [75] with default parameters. To perform Principal Component Analysis (PCA), Beagle4.1 [79] was used to impute the unphased genotypes. All the imputed and identified genotypes at SNP loci were pooled and finalized using PLINK [77] and iTools [80], which were then subjected to PCA analysis using a software GCTA [81]. The population was clustered using ADMIXTURE software [35] with K (the expected number of clusters) increasing from 2 to 9. The K value with the minimum cross-validation error was eventually selected.

Genome-wide linkage disequilibrium (LD) was calculated for populations of different groups using Haploview (http://www.broadinstitute.org/haploview/haploview) in windows of 2,000 kb. Briefly, the correlation coefficient (r^*2*^) between SNP pairs in a non-overlapping sliding 1 kb bin was calculated and then averaged within bins.

We identified candidate regions under selection by comparing polymorphism levels, measured by *ROD* as well as *F*_*ST*_, between “Gilo”, “Shum” and “*Solanum anguivi*” groups. *ROD* was calculated following the formula of: *ROD* = 1 - π_cul_/π_wild_, where π_cul_ and π_wild_ denote the nucleotide diversity within the cultivated and wild populations, respectively. *F*_*ST*_ measurement was calculated according to the formula: *F*_*ST*_ = (π_between_-π_within_)/ π_between._ where π_between_ and π_within_ represent the average number of pairwise differences between two individuals sampled from different or the same population.

### Construction of pan- and core-genome

To build a gene set including *S. atehiopicum* genes as many as possible, we assembled the contigs of all the re-sequenced 65 accessions individually using SOAPdenovo2 [38]. The assembled contigs from each group (“Gilo”, “Shum” and “*S. anguivi*”) were then merged together. CD-HIT-EST [39] was used to eliminate the redundancy and generate the final dataset of pan-genomes for each group. Similarly, all these contigs were merged into a pan-genome of *S. aethiopicum*. Gene models were predicted from these contigs as described above and their functions were also annotated.

## Abbreviations

4DTV: four-fold degenerative third-codon transversion
BUSCO: Benchmarking Universal Single-Copy Orthologs
CEG: core embryophyta gene
CV: cross-validation
GATK: Genome Analysis Toolkit
LTR: long terminal repeat
LINE: long interspersed element
LD: Linkage disequilibrium
MYA: million years ago
PSMC: pairwise sequential Markovian coalescent model
PCA: principal-component analysis
SINE: short interspersed element
TE: transposable elements
WGD: whole genome duplication
WGS: whole-genome shotgun

## Availability of supporting data

The raw sequence data from our genome project, including assembly and annotation of *S. aethiopicum* genome, as well as the re-sequencing data from 65 samples, were deposited in the CNGB Nucleotide Sequence Archive database (CNSA: https://db.cngb.org/cnsa.) under project accession number CNP0000317. All supplementary figures and tables are provided as Additional Files.

## Additional files

Supplementary Tables-1.docx

Supplementary Tables-2.rar

Supplementary Figures.docx

## Authors contributions

X. X., A. D., X. L., J. W., and H. Y. conceived the project; S. C. and H. L. managed and supervised these works; B. S. and Y. F. managed the samples in BGI; B. S. and Y. F assembled, Y. F and Y. S. annotated the genome. H. L. and S. P. constructed DNA libraries and sequenced the genome. Y. S and B. S. performed the analysis of gene families, LTR evolution, and transcriptomic data; Y. F., B. S., and Y. S. analyzed the re-sequencing data; Y. S., Y. F. and B. S. collected dataset required for the analysis in this study. B. S, X. L., Y. S., D. O., and Y. F. wrote and revised the manuscript.

