## Supplemental Tables for "Draft genome sequence of the *Solanum aethiopicum* provides insights into disease resistance, drought tolerance and the evolution of the genome"

**Supplementary Table S1** Summary of the library types and data generated in this study

| Insert size  (bp) | Raw data | | Clean data | |
| --- | --- | --- | --- | --- |
|  | Total data  (Gb) | Sequence depth (X) | Total data  (Gb) | Sequence depth  (X) |
| 250 | 54.80 | 46.90 | 47.68 | 40.81 |
| 500 | 37.84 | 32.39 | 31.95 | 27.34 |
| 2000 | 33.69 | 28.83 | 15.38 | 13.16 |
| 6000 | 40.81 | 34.92 | 16.03 | 13.72 |
| 10000 | 43.29 | 37.05 | 6.31 | 5.40 |
| 20000 | 32.18 | 27.55 | 10.47 | 8.97 |
| Total | 242.61 | 207.64 | 127.83 | 109.40 |

**Supplementary Table S2**	Statistics of the *S. aethiopicum* assembly

|  | Scaffold | | Scaftig | |
| --- | --- | --- | --- | --- |
|  | Length(bp) | Number | Length(bp) | Number |
| Max length | 2,941,814 | 366,216 |  |  |
| N90 | 59,241 | 2,488 | 3,091 | 47,049 |
| N80 | 174,853 | 1,533 | 8,181 | 29,145 |
| N70 | 284,610 | 1,072 | 13,361 | 20,239 |
| N60 | 385,168 | 763 | 18,957 | 14,355 |
| N50 | 516,145 | 536 | 25,233 | 10,078 |
| N40 | 680,480 | 364 | 32,488 | 6,807 |
| N30 | 879,386 | 232 | 42,235 | 4,275 |
| N20 | 1,138,385 | 127 | 55,207 | 2,326 |
| N10 | 1,625,202 | 50 | 77,675 | 879 |
| Total length | 1,028,783,629 | -- | 936,403,134 | -- |
| num>=100bp | 162,186 | 231,821 | -- | -- |
| num>=2kb | 8,839 | 55,524 | -- | -- |
| Rate if gap | 8.98% | -- | -- | -- |

**Supplementary Table S3** Comparison of the genomic characteristics in different genomes

| Species | *S. aethiopicum* | pepper | Tomato | Potato |
| --- | --- | --- | --- | --- |
| Assembled genome size (Mb) | 1,028 | 3,349 | 760 | 727 |
| GC count（%） | 33.13 | 34.90 | 34.00 | 34.80 |
| Repeat rate (%) | 78.90 | 80.90 | 63.20 | 62.20 |
| Gene number | 34,906 | 35,336 | 34,727 | 35,004 |

**Supplementary Table S4** Statistics of repeat annotation, transposable elements

|  | Type | %in genome | Length(bp) |
| --- | --- | --- | --- |
| **Type I: Retrotransposon elements** | |  |  |
|  | SINE | 0.42 | 4,420,139 |
|  | LINE | 5.06 | 52,062,362 |
|  | LTR | 69.94 | 719,631,011 |
|  | Other | 0.003 | 33,607 |
| **Type II: DNA transposon** | |  |  |
|  | DNA | 2.87 | 29,601,166 |
| **Type III: Tandem repeats** | |  |  |
|  | Satellite | 0.0007 | 7,614 |
|  | Simple repeat | 0.15 | 1,562,537 |
| **unknow** |  | 0.42 | 4,390,358 |
| **Others** |  | 0.0003 | 3,648 |
| **Total repeat** |  | 78.9 | 811,712,442 |

**Supplementary Table S5**	Statistics of gene model in different species

| Species | # of genes | Average gene lenth (bp) | Average CDS length (bp) | Average exon number | Average exon length (bp) | Average intron length (bp) |
| --- | --- | --- | --- | --- | --- | --- |
| *S.aethiopicum* | 34,906 | 3038.08 | 1104.26 | 4.15 | 265.82 | 613.1 |
| *S.lycopersicum* | 34,727 | 2942.39 | 1035.86 | 4.53 | 228.78 | 540.42 |
| *S.tuberosum* | 38,492 | 2476.45 | 928.01 | 3.49 | 265.58 | 620.79 |
| *C.annuum* | 45,131 | 6439.84 | 1275.46 | 5.58 | 228.5 | 1127.15 |
| *N.sylvestris* | 33,709 | 3956.67 | 1172.7 | 4.73 | 248.04 | 746.8 |
| *A.thaliana* | 26,637 | 1909.57 | 1242.78 | 5.23 | 237.5 | 157.54 |

**Supplementary Table S6**	Evaluation of predicted gene models using 1,440 CEGs

| Type | Number | Percentage (%) |
| --- | --- | --- |
| Complete BUSCOs | 1158 | 80.4% |
| Complete and single-copy BUSCOs | 1120 | 77.8% |
| Complete and duplicated BUSCOs | 38 | 2.6% |
| Fragmented BUSCOs | 111 | 7.7% |
| Missing BUSCOs | 171 | 11.9% |
| Total BUSCO groups searched | 1440 |  |

**Supplementary Table S7**	Statistics of predicted ncRNA

| Type |  | Number | Average length(bp) | Total length(bp) | % of genome |
| --- | --- | --- | --- | --- | --- |
| **miRNA** |  | 128 | 121.7031 | 15578 | 0.0015 |
| **tRNA** |  | 960 | 75.1865 | 72179 | 0.0070 |
| **rRNA** | rRNA | 1185 | 129.2852 | 153203 | 0.0149 |
|  | 18S | 157 | 347.8981 | 54620 | 0.0053 |
|  | 28S | 126 | 119.0556 | 15001 | 0.0015 |
|  | 5.8S | 32 | 127.1250 | 4068 | 0.0004 |
|  | 5S | 870 | 91.3954 | 79514 | 0.0077 |
| **snRNA** | snRNA | 503 | 117.1133 | 58908 | 0.0057 |
|  | CD-box | 223 | 100.4395 | 22398 | 0.0022 |
|  | HACA-box | 51 | 123.8235 | 6315 | 0.0006 |
|  | splicing | 229 | 131.8559 | 30195 | 0.0029 |

**Supplementary Table S8**	Result of function annotation

| Database | Number | Percentage (%) |
| --- | --- | --- |
| Nr Annotated | 30,842 | 88.36 |
| Swissprot Annotated | 20,932 | 59.97 |
| KEGG Annotated | 20,077 | 57.52 |
| COG Annotated | 9,894 | 28.34 |
| TrEMBL Annotated | 31,099 | 89.09 |
| Interpro Annotated | 26,319 | 75.4 |
| GO Annotated | 5,101 | 14.61 |
| Total functional gene | 31,863 | 91.28 |

**Supplementary Table S9**	Statistics of gene families

| Species | Genes number | Genes in families | Unclustered genes | Family number | Unique families | Average genes per family |
| --- | --- | --- | --- | --- | --- | --- |
| *S. lycopersicum* | 34,727 | 26,804 | 7,923 | 18,346 | 429 | 1.46 |
| *S. aethiopicum* | 34,906 | 25,751 | 9,155 | 19,310 | 465 | 1.33 |
| *S. tuberosum* | 38,492 | 31,692 | 6,800 | 18,023 | 610 | 1.76 |
| *C. annuum* | 35,884 | 27,840 | 8,044 | 16,177 | 792 | 1.72 |
| *S. melongena* | 85,446 | 72,446 | 13,000 | 19,898 | 2,449 | 3.64 |

**Supplementary Table S10** Common shared gene families between *S. aethiopicum* and other species

| Speices | # of common shared gene families with *S. aethiopicum* |
| --- | --- |
| *S. melongena* | 15,723 |
| *S. lycopersicum* | 15,379 |
| *S. tuberosum* | 14,820 |
| *C. annuum* | 13,461 |

**Supplementary Table S15** Number of annotated NLR genes in *S. aethiopicum*, *S. melongena* and other species

| Speices | Annotated NLR genes | Retrogene (%) |
| --- | --- | --- |
| *C. annuum** | 883 | 123 (13.9) |
| *S. aethiopicum* | 447 | 62 (13.8%) |
| *S. melongena* | 251 | 33 (13.1%) |
| Tomato* | 267 | 21 (7.8%) |
| Potato* | 443 | 81 (18.2%) |

*** According to Kim et al (2017).

**Supplementary Table S20** Pan genome annotation

| Group | Anguivi | Gilo | Shum | All |
| --- | --- | --- | --- | --- |
| gene number | 17662 | 33194 | 51351 | 41626 |
| average gene length | 1704.85 | 1650.43 | 1624.21 | 1641.12 |
| average CDS length | 972.08 | 935.51 | 921.01 | 929.81 |
| average exon length | 285.07 | 281.78 | 285.85 | 288.37 |
| average exon number | 3.41 | 3.32 | 3.22 | 3.22 |
| average intron length | 304.06 | 308.16 | 316.48 | 319.79 |
| average intron number | 2.41 | 2.32 | 2.22 | 2.22 |
