## Supplementary Figures for "Draft genome sequence of the *Solanum aethiopicum* provides insights into disease resistance, drought tolerance and the evolution of the genome"

**Figure S1** 17-mer analysis for estimating the *S. aethiopicum* genome size. The peak of distribution is about 38X coverage, then the genome size can be estimated as ~1.16 Gb (Genome Size=K-mer number/Peak depth). The small peak at 1/2 the peak depth shows the high heterozygous rate of the genome.

Kmer frequency

| Depth(X) |  | 1 |
| --- | --- | --- |
| 0 | 36.152 | 1 |
| 1 | 11.2507 |  |
| 2 | 3.57777 |  |
| 3 | 1.53991 |  |
| 4 | 0.655209 |  |
| 5 | 0.360372 |  |
| 6 | 0.215543 |  |
| 7 | 0.153361 |  |
| 8 | 0.121022 |  |
| 9 | 0.106754 |  |
| 10 | 0.101471 |  |
| 11 | 0.102852 |  |
| 12 | 0.108136 |  |
| 13 | 0.116937 |  |
| 14 | 0.128249 |  |
| 15 | 0.142191 |  |
| 16 | 0.159136 |  |
| 17 | 0.179203 |  |
| 18 | 0.203899 |  |
| 19 | 0.233695 |  |
| 20 | 0.269359 |  |
| 21 | 0.312211 |  |
| 22 | 0.365179 |  |
| 23 | 0.427344 |  |
| 24 | 0.500293 |  |
| 25 | 0.584316 |  |
| 26 | 0.677365 |  |
| 27 | 0.779731 |  |
| 28 | 0.888891 |  |
| 29 | 1.00248 |  |
| 30 | 1.11389 |  |
| 31 | 1.22113 |  |
| 32 | 1.31929 |  |
| 33 | 1.40376 |  |
| 34 | 1.47127 |  |
| 35 | 1.5199 |  |
| 36 | 1.54653 |  |
| 37 | 1.55052 |  |
| 38 | 1.53217 |  |
| 39 | 1.49283 |  |
| 40 | 1.43541 |  |
| 41 | 1.36142 |  |
| 42 | 1.27446 |  |
| 43 | 1.17834 |  |
| 44 | 1.0765 |  |
| 45 | 0.972762 |  |
| 46 | 0.869477 |  |
| 47 | 0.770185 |  |
| 48 | 0.675485 |  |
| 49 | 0.588056 |  |
| 50 | 0.509212 |  |
| 51 | 0.438546 |  |
| 52 | 0.376726 |  |
| 53 | 0.324093 |  |
| 54 | 0.280227 |  |
| 55 | 0.243739 |  |
| 56 | 0.214479 |  |
| 57 | 0.191653 |  |
| 58 | 0.1746 |  |
| 59 | 0.162567 |  |
| 60 | 0.15452 |  |
| 61 | 0.14958 |  |
| 62 | 0.147803 |  |
| 63 | 0.148383 |  |
| 64 | 0.150527 |  |
| 65 | 0.153959 |  |
| 66 | 0.15793 |  |
| 67 | 0.162705 |  |
| 68 | 0.167526 |  |
| 69 | 0.172209 |  |
| 70 | 0.176345 |  |
| 71 | 0.18041 |  |
| 72 | 0.183149 |  |
| 73 | 0.185268 |  |
| 74 | 0.186633 |  |
| 75 | 0.186662 |  |
| 76 | 0.185971 |  |
| 77 | 0.184381 |  |
| 78 | 0.181962 |  |
| 79 | 0.17891 |  |
| 80 | 0.173968 |  |
| 81 | 0.169114 |  |
| 82 | 0.162913 |  |
| 83 | 0.157043 |  |
| 84 | 0.150318 |  |
| 85 | 0.143468 |  |
| 86 | 0.136267 |  |
| 87 | 0.12922 |  |
| 88 | 0.122233 |  |
| 89 | 0.115292 |  |
| 90 | 0.108599 |  |
| 91 | 0.102285 |  |
| 92 | 0.0964161 |  |
| 93 | 0.091044 |  |
| 94 | 0.0859105 |  |
| 95 | 0.0813188 |  |
| 96 | 0.0773257 |  |
| 97 | 0.0738172 |  |
| 98 | 0.0704916 |  |
| 99 | 0.0679766 |  |
| 100 | 0.0655161 |  |
| 101 | 0.0637682 |  |
| 102 | 0.0621513 |  |
| 103 | 0.0610571 |  |
| 104 | 0.059938 |  |
| 105 | 0.0592241 |  |
| 106 | 0.0583745 |  |
| 107 | 0.0580807 |  |
| 108 | 0.0575489 |  |
| 109 | 0.0573233 |  |
| 110 | 0.0571313 |  |
| 111 | 0.0568492 |  |
| 112 | 0.0563858 |  |
| 113 | 0.0560981 |  |
| 114 | 0.0556413 |  |
| 115 | 0.0551103 |  |
| 116 | 0.0545618 |  |
| 117 | 0.0539456 |  |
| 118 | 0.0531887 |  |
| 119 | 0.0522982 |  |
| 120 | 0.0514844 |  |
| 121 | 0.050579 |  |
| 122 | 0.0495353 |  |
| 123 | 0.0485531 |  |
| 124 | 0.0473213 |  |
| 125 | 0.0460799 |  |
| 126 | 0.044985 |  |
| 127 | 0.0437902 |  |
| 128 | 0.0424925 |  |
| 129 | 0.0413714 |  |
| 130 | 0.0403778 |  |
| 131 | 0.0392529 |  |
| 132 | 0.0379846 |  |
| 133 | 0.037047 |  |
| 134 | 0.036022 |  |
| 135 | 0.0348742 |  |
| 136 | 0.0341046 |  |
| 137 | 0.0333504 |  |
| 138 | 0.032533 |  |
| 139 | 0.0318428 |  |
| 140 | 0.0310837 |  |
| 141 | 0.0305306 |  |
| 142 | 0.0300534 |  |
| 143 | 0.029496 |  |
| 144 | 0.0289525 |  |
| 145 | 0.0285839 |  |
| 146 | 0.0283087 |  |
| 147 | 0.0278562 |  |
| 148 | 0.0274622 |  |
| 149 | 0.0271337 |  |
| 150 | 0.026835 |  |
| 151 | 0.0266235 |  |
| 152 | 0.0264273 |  |
| 153 | 0.0261258 |  |
| 154 | 0.0258133 |  |
| 155 | 0.0254596 |  |
| 156 | 0.02515 |  |
| 157 | 0.0248569 |  |
| 158 | 0.0245897 |  |
| 159 | 0.0243113 |  |
| 160 | 0.023971 |  |
| 161 | 0.0236485 |  |
| 162 | 0.0232738 |  |
| 163 | 0.0228818 |  |
| 164 | 0.0225384 |  |
| 165 | 0.0222866 |  |
| 166 | 0.0219049 |  |
| 167 | 0.0215074 |  |
| 168 | 0.0212041 |  |
| 169 | 0.020855 |  |
| 170 | 0.0204783 |  |
| 171 | 0.0201405 |  |
| 172 | 0.0197279 |  |
| 173 | 0.0194149 |  |
| 174 | 0.0191704 |  |
| 175 | 0.0188213 |  |
| 176 | 0.0185517 |  |
| 177 | 0.0182596 |  |
| 178 | 0.0180492 |  |
| 179 | 0.0176874 |  |
| 180 | 0.0173898 |  |
| 181 | 0.0172616 |  |
| 182 | 0.0168935 |  |
| 183 | 0.0166304 |  |
| 184 | 0.0165227 |  |
| 185 | 0.0163146 |  |
| 186 | 0.0161035 |  |
| 187 | 0.0159652 |  |
| 188 | 0.015786 |  |
| 189 | 0.0155178 |  |
| 190 | 0.0153034 |  |
| 191 | 0.0151989 |  |
| 192 | 0.0150199 |  |
| 193 | 0.0149641 |  |
| 194 | 0.0147906 |  |
| 195 | 0.0145083 |  |
| 196 | 0.0143881 |  |
| 197 | 0.0142059 |  |
| 198 | 0.0141454 |  |
| 199 | 0.0139254 |  |

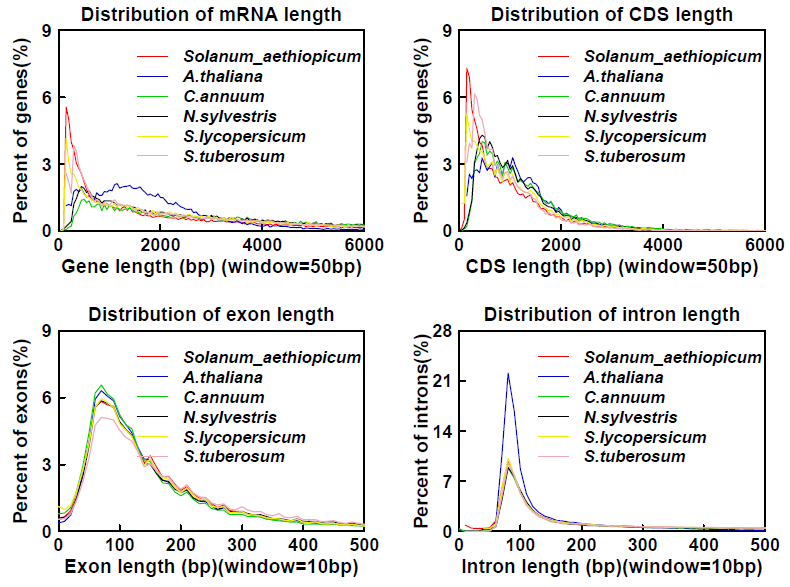

**Figure S2** Distributions of the gene model for four categories in the relative species.

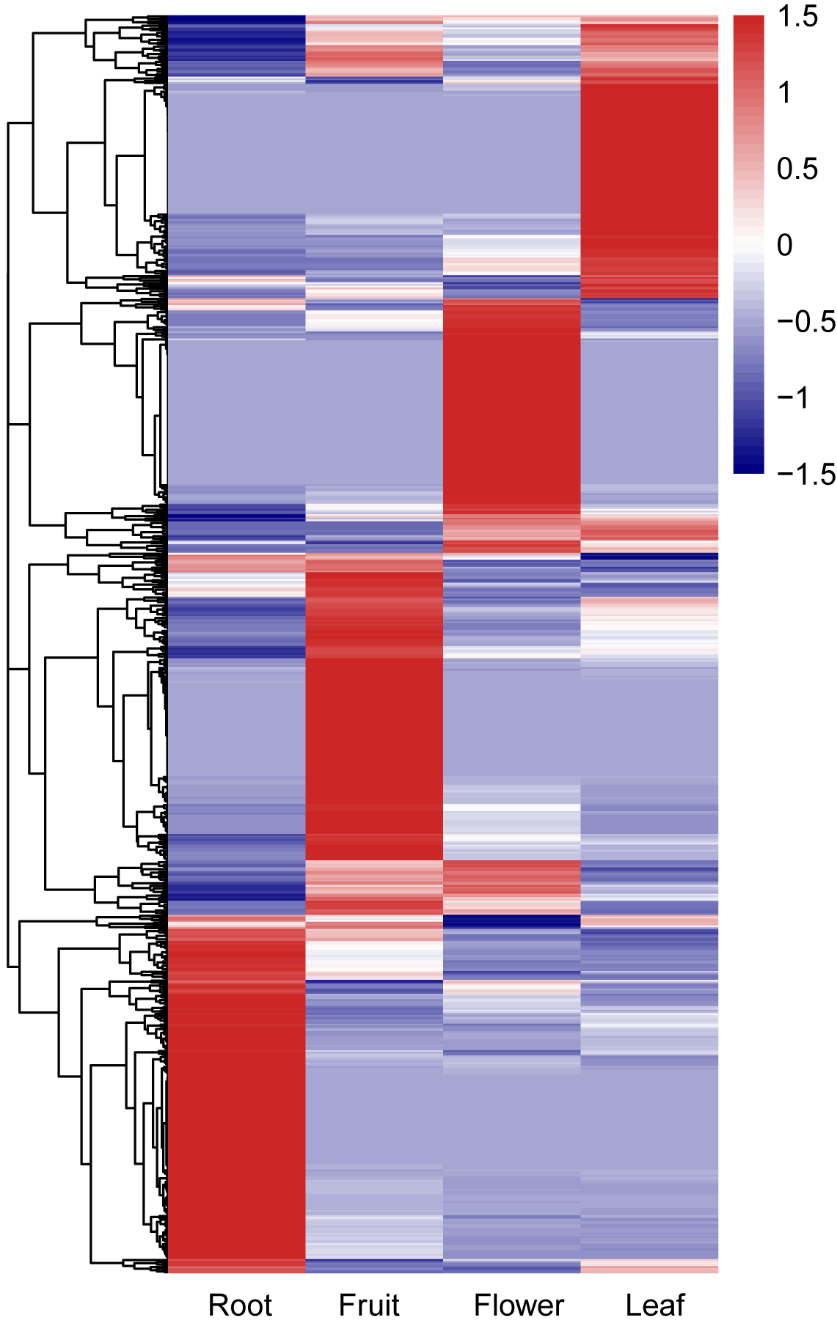

**Figure S3** Gene expression of LTR-captured gene in different tissues

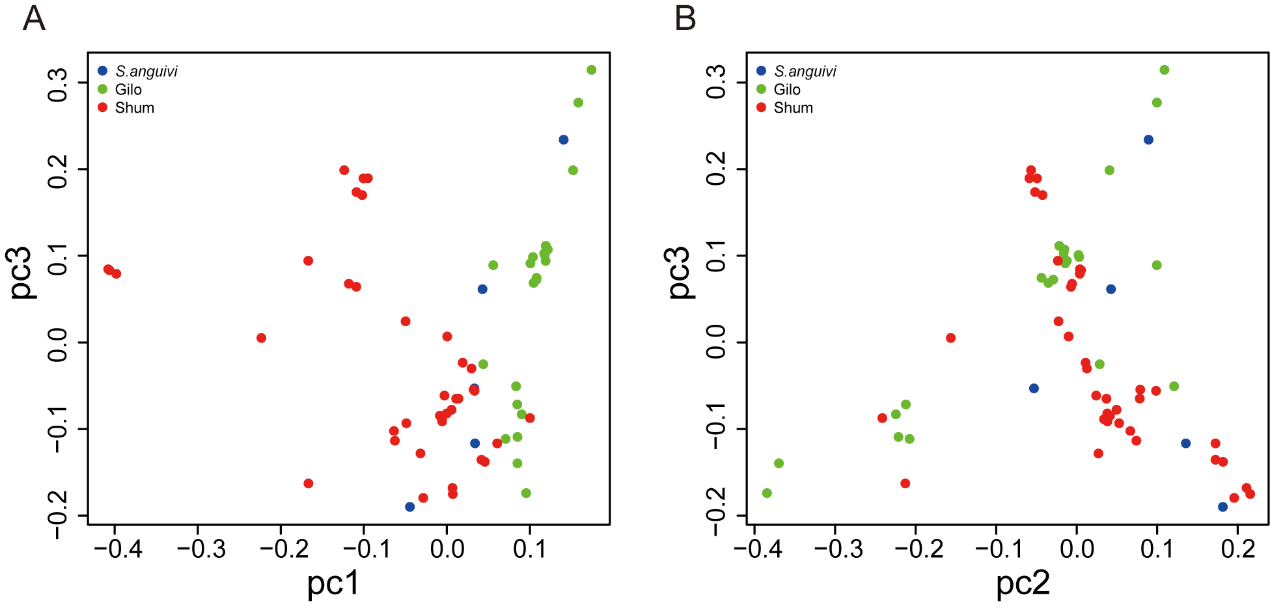

**Figure S4** Principal-component analysis
